# Arousal increases locus coeruleus blood flow, salience-related brain responses, and modulates negative-valence attentional biases

**DOI:** 10.1101/2025.10.07.680973

**Authors:** Andy J. Kim, Chenyang Zhao, Fanhua Guo, Ioannis Pappas, Martin J. Dahl, Heidi I.L. Jacobs, Danny J.J. Wang, Mara Mather

**Affiliations:** Leonard Davis School of Gerontology, University of Southern California, Los Angeles, CA, USA; Laboratory of FMRI Technology, Mark & Mary Stevens Neuroimaging and Informatics Institute, Keck School of Medicine, University of Southern California, Los Angeles, CA, USA; Center for Lifespan Psychology, Max Planck Institute for Human Development, Berlin, Germany; Max Planck UCL Centre for Computational Psychiatry and Ageing Research; Athinoula A. Martinos Center for Biomedical Imaging, Department of Radiology, Massachusetts General Hospital, Harvard Medical School, Boston, Massachusetts, USA; Department of Psychology, University of Southern California, Los Angeles, CA, USA; Department of Biomedical Engineering, University of Southern California, Los Angeles, CA, USA

**Keywords:** locus coeruleus, valence, arousal, fMRI, emotion, attention

## Abstract

The amygdala helps prioritize emotional over neutral information. However, it responds similarly to positive and negative stimuli, and so is unlikely to be the source of valence-specific effects within affective networks. We hypothesized that the locus coeruleus (LC) is a key contributor to negative biases in attention. Using ultra-high field 7T magnetic resonance imaging, we tested how arousal modulates processing of emotional faces during an oddball task in twenty-two young adults during two separate sessions. Arousal induced by isometric handgrip increased LC cerebral blood flow and amplified brain responses to target and angry faces, but not to happy faces. The amygdala exhibited valence-general responses that were not modulated by arousal. LC connectivity with the default mode network decreased during processing target and angry faces, and arousal further modulated responses in the salience network and visual cortex. Behaviorally, arousal enhanced recognition of angry faces only when allocating attentional resources and memory performance was linked to left LC brain activity. These findings highlight the LC as a key structure through which arousal shapes valence processing, biases attention, and informs mechanisms related to affective disorders.

## Introduction

Emotion is conceptualized along two core dimensions: arousal (reflecting physiological activation) and valence (reflecting positive or negative affect). In the brain, the amygdala contributes to these dimensions of emotion by mediating the influence of arousal on cognitive processes (LaBar & Cabeza, 2006) and encoding both positive and negative valence (Paton et al., 2006). In addition, the locus coeruleus (LC), a small brainstem nucleus (∼75 mm^3^) that serves as the brain’s principal source of the neuromodulator noradrenaline, also responds to both positive and negative stimuli, and helps coordinate global arousal regulation and vigilance states that shape cognition, including attention (Thiele & Bellgrove, 2018). Noradrenergic fibers from the LC innervate the amygdala via the ansa peduncularis-ventral amygdaloid bundle (Fallon et al., 1978; Schwarz et al., 2015), and the amygdala sends reciprocal projections to the LC (Gu et al., 2020), creating a bidirectional amygdala-LC circuit. In rodents, pharmacological manipulations of noradrenaline and optogenetic stimulation of LC projections modulate downstream amygdala responses (McCall et al., 2017; Tully & Bolshakov, 2010; Uematsu et al., 2017), and chemogenetic stimulation of the LC increases freezing behavior and suppresses neuronal activation in the ventromedial prefrontal cortex through *β-*noradrenergic receptor signaling in the basolateral amygdala (Bayer et al., 2025). Furthermore, in humans, the LC interacts with the amygdala to facilitate the consolidation of emotionally salient experiences (Jacobs et al., 2020). Functional coupling between the LC and amygdala predicts individual differences in stress-related anxiety symptoms in real-world contexts (Grueschow et al., 2021), and increases during the encoding of novel face-name associations (Prokopiou et al., 2022). Together, these findings establish the amygdala-LC circuit as a core mechanism through which arousal and valence actively shape affective behavior.

Meta-analyses of fMRI and PET studies indicate that the amygdala exhibits increased activation in response to both positive- and negative-valence arousing stimuli compared with non-arousing stimuli (Costafreda et al., 2008; Sergerie et al., 2008), across both visual and auditory domains (Lin et al., 2020). These findings suggest that the amygdala primarily encodes arousal intensity, while its role in valence processing emerges through broader connectivity with cortical and subcortical circuits that assign positive or negative value (Pessoa & Adolphs, 2010). Given that the amygdala responds in a valence-general manner, the prevalence of robust valence-specific effects in the affective attention literature remains an unresolved mechanistic question. Emotional stimuli are particularly prioritized by the attention system and elicit increased automatic orienting compared with neutral stimuli (Carretié, 2014; Vuilleumier, 2005), a bias that likely evolved across species to promote survival (Baumeister et al., 2001; Rozin & Royzman, 2001). Foundational models of LC-noradrenaline system (LC-NA) function characterize the LC as a regulator of adaptive gain and network resetting in response to salient events, with its activity primarily driven by arousal intensity (Aston-Jones & Cohen, 2005; Bouret & Sara, 2005). Through its widespread cortical and subcortical projections, the LC releases noradrenaline during arousing states, biasing attentional selection toward salient information (Markovic et al., 2014; Mather et al., 2016). Under arousal, LC activation has shown to selectively enhance memory and attention for both prioritized and salient information (Clewett et al., 2018; Lee et al., 2018). Furthermore, studies in rodents show that LC optogenetic activation encode both positive and negative valence odors through distinct firing patterns and adrenoreceptors, in which conditioning could be prevented with pharmacological blockades in the basolateral amygdala (Ghosh et al., 2021; Omoluabi et al., 2022). These findings position the LC as a central driver of how arousal shapes emotional attention and incorporates valence, either through dynamic interactions with the amygdala or as an independent system.

In this study, we hypothesized that the LC, rather than the amygdala, primarily modulates stimulus valence within arousal networks. Unfortunately, neuroimaging of the LC poses significant technical challenges due to its small size and deep location within the brainstem (Forstmann et al., 2016). Previous work have capitalized on the structural properties of the LC to measure changes in MRI contrast (Bueichekú et al., 2024; Dahl et al., 2023), but studies measuring task-related LC brain responses remain scarce (Berger et al., 2023; Hall et al., 2024; Jacobs et al., 2020; Krebs et al., 2018; Ludwig et al., 2024; Murphy et al., 2014). In a visual oddball task, Krebs et al. (2018) demonstrated that LC responses increased significantly to novel negative and novel neutral stimuli compared with frequent stimuli. Therefore, we used ultra-high field (7T) neuroimaging to improve brainstem signal-to-noise ratio and modified the emotional oddball paradigm to include novel happy faces in addition to novel angry faces, enabling investigation of valence-specific LC responses. To manipulate arousal during scanning, we implemented an isometric handgrip paradigm which increases LC-NA system activity as indexed by proxy measures such as pupil diameter, salivary amylase, and heart rate (Bachman et al., 2023; Mather et al., 2020; Nielsen & Mather, 2015). We investigated how arousal modulates functional LC responses, its interactions with attention networks, and recognition memory for emotional faces. Finally, we present a novel high-resolution (1 mm isotropic) whole-brain perfusing imaging approach using an ultra-high field MRI protocol. By combining an advanced pseudo-continuous arterial spin labeling (pCASL) pulse sequence with a dynamic compressed sensing reconstruction algorithm (Jung et al., 2009; Lustig et al., 2007; Wang et al., 2022), we non-invasively indexed brainstem metabolic activity to determine whether the arousal manipulation exerted specific or broad effects within the brainstem and the amygdala.

## Methods

### Participants and Study Design

Twenty-two participants completed two separate sessions of the experiment (mean age: 24.8 years old, std. dev = 4.6, range: 18-32; female = 16; see Figure 1). All study protocols were approved by the University of Southern California Institutional Review Board (UP-24-00514) and written informed consent was obtained from all participants prior to participation. In addition, nine participants completed additional arterial spin labeling (ASL) scans. Participants completed two experimental sessions on separate days (mean = 2.9 days apart) scheduled at approximately the same time of day. Each session lasted two hours. At the beginning of each session, participants completed the task portion of the experiment in the scanner. This portion consisted of a T1-weighted MP2RAGE anatomical scan, followed by a block consisting of one magnetization transfer (MT) structural scan targeting the brainstem and two runs of the oddball task. This block was repeated, resulting in a second MT scan and two additional task runs. During one of the two MT scans, participants performed an isometric handgrip task, alternating between hands when fatigued, and thus consistently squeezing during the entire duration of the scan. To counterbalance task order, half of the participants performed the handgrip task during the first MT scan block, while the other half completed it during the second MT scan block. The block with and without the handgrip intervention is referred to as the control or arousal condition, respectively. After the task portion, participants exited the scanner to complete a recognition memory task and a brief break. Participants then returned to the scanner for ASL scans which included three conditions: baseline (pre-handgrip), handgrip, and rest (post-handgrip). As in the earlier structural MT scan, participants squeezed the handgrip continuously during the handgrip ASL scan. The procedure on session two was identical to session one except that the order of the handgrip condition was reversed to achieve full counterbalancing; participants who completed the handgrip during the first block on Day 1 performed it during the second block on day 2, and *vice versa*.

**Figure 1.**
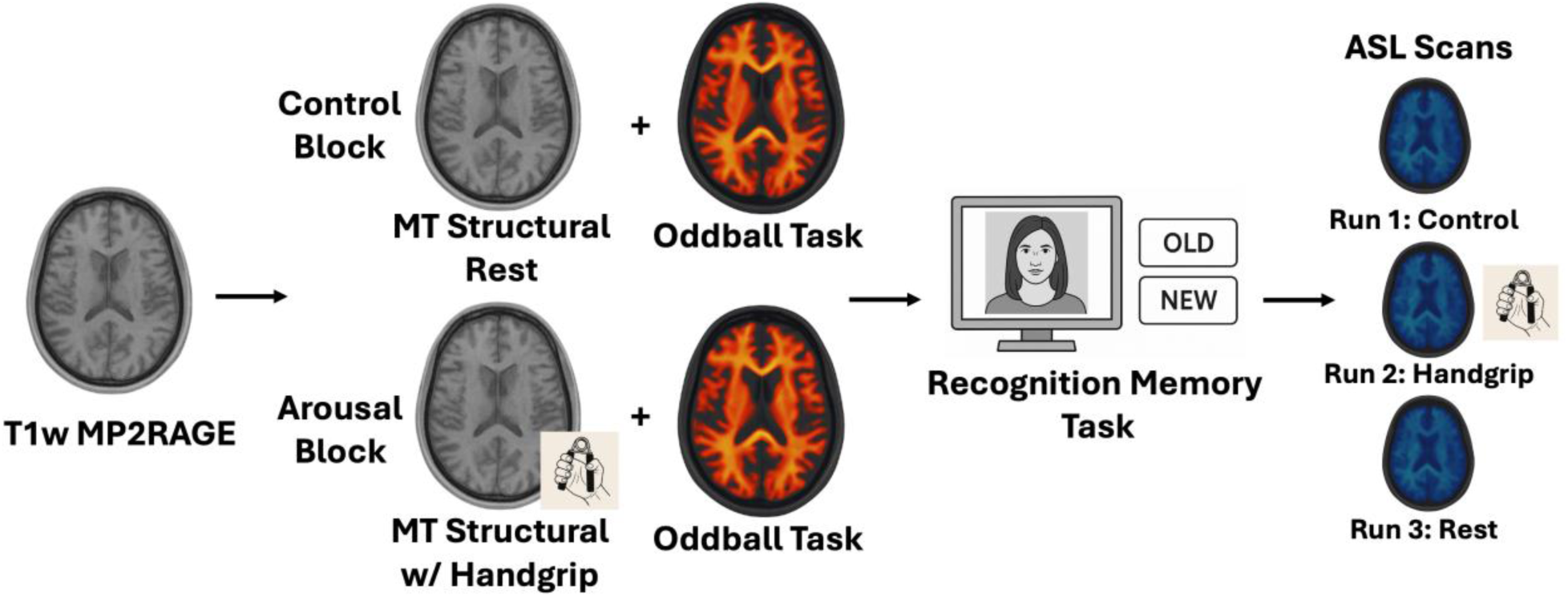
Experiment Procedure. Following the structural anatomical scan, participants completed a control or arousal block consisting of a magnetization transfer (MT) structural scan to measure LC MRI contrast and two runs of the oddball task. The order of block conditions was counterbalanced across participants per session and across both sessions (e.g., if completing the control block first on Session 1, the arousal block was completed first on Session 2). Next, participants exited the scanner and completed a recognition memory task. Finally, nine participants additionally completed three arterial spin labeling (ASL) scans: a pre-handgrip period (*control*), an arousal condition while squeezing the isometric handgrip (*handgrip*), and a post-handgrip period (*rest*). The emotional oddball task included standard (repetitive; 70% of trials), target (10% of trials), novel positive (10% of trials), and novel negative (10% of trials) faces. The target face was the only older adult, and all other faces were young adults. For the novel emotional faces, gender and race were counterbalanced (50% female, 50% male; 33% White, 33% Black, 33% Asian).

### Emotional oddball task procedures and stimuli

Participants completed four runs of the emotional oddball task each session (two runs each block; see Figure 1). The experimental design was modified from a previous emotional oddball paradigm measuring LC brain responses (Krebs et al., 2018). Each run comprised of 150 total trials: 105 repetitions of a familiar neutral face (standard face; 70%), 15 repetitions of a familiar older adult face (target face; 10%), 15 novel happy faces (positive valence; 10%), and 15 angry faces (negative valence; 10%). All stimuli were color photographs selected from the following three sources: FACES database (Ebner et al., 2010), Chicago Face Database (Ma et al., 2015), and Chinese Face dataset (Han et al., 2023). All faces were of young adults except the target older adult. The novel valence faces were balanced for gender (50% male, 50% female) and race (approximately 33% White, 33% Black, 33% Asian). Trials were randomized each run. On each trial, a face stimulus appeared centrally at fixation for 800 ms, followed by a jittered inter-trial interval (ITI) consisting of a central fixation cross presented for 1.6-2.4 seconds (1.6, 1.8, 2.0, 2.2, or 2.4 seconds, equally often). A fixation cross was continuously displayed throughout the task and overlaid on all face stimuli. Participants were instructed to maintain fixation and silently count the number of target face presentations. At the end of each run, participants verbally reported their target face count to the experimenter to enforce task adherence.

### Recognition memory task

Following the task portion, participants exited the scanner and completed a recognition memory task. This test included all novel emotional faces previously presented during the functional task scans, along with an additional set of thirty happy and angry faces each that had not been shown. Participants viewed each face individually and indicated whether they had seen that face during the task (Yes/No recognition). Outcomes were analyzed separately for faces presented in the control and handgrip blocks. For each subject and condition, hit rate was calculated as the proportion of correctly identified items (hits; 60 instances per valence per condition) and the false alarm rate as the proportion of new items incorrectly judged as old (false alarms; 30 instances per valence). Log-linear corrections were applied to avoid infinite *z*-scores (Hautus, 1995).

### Magnetic resonance imaging (MRI) sequences

MRI data were collected using a 7T Terra system (Siemens Medical System, Erlangen, Germany). For the structural and functional scans, data were collected with a 1Tx/32Rx head coil (Nova Medical, Cambridge, MA, USA). A sagittal T1-weighted magnetization-prepared rapid acquisition with gradient echo sequence (T1w-MP2RAGE) sequence was collected to provide high-resolution anatomical images for co-registration with the following acquisition parameters: matrix size = 231 x 256 x 192, voxel size = 1.1 x 1.1 x 1.0 mm, repetition time = 4300 ms, echo time = 1.84 ms, inversion times = 840/2370 ms, flip angles = 5°/6 °, and acquisition time = 5:24 minutes. Axial magnetization transfer structural scans (Priovoulos et al., 2018) optimized for brainstem imaging were acquired in a slab FOV (60 slices, 192 mm, 0.5 mm slice thickness) covering the brainstem with the following acquisition parameters: matrix size = 263 x 350 x 350, voxel size = 0.4 x 0.4 x 0.5 mm, repetition time = 400 ms, echo time = 2.55 ms, flip angle = 8°, and acquisition time = 8:13 minutes. Finally, T2*-weighted gradient-echo echo-planar imaging (EPI) sequences were acquired to measure blood-oxygen-level-dependent (BOLD) signals during task performance with the following acquisition parameters: matrix size = 134 x 134 mm, 84 interleaved slices, 219 measurements, voxel size = 1.5 mm^3^ isotropic, repetition time = 2000 ms, echo time = 27.40 ms, flip angle = 45°, multiband acceleration factor = 3, in-plane parallel imaging: GRAPPA with acceleration factor = 2, and acquisition time = 7:46 minutes.

For arterial spin labeling (ASL) scans, data were collected with an 8Tx/32Rx head coil (Nova Medical) on the investigational pTx part of the 7T Terra system. Imaging was conducted using whole-brain 3D pseudo-continuous arterial spin labeling (3D pCASL) with 1 mm isotropic resolution (Wang et al., 2022). Data were acquired using the rotated golden-angle stack-of-spirals (rGA-SOS) sampling scheme (Munsch et al., 2020) with the following parameters: field of view (FOV) = 256 x 256 x 144 mm^3^, one spiral shot per TR, and 144 kHz partitions filled per TR (Guo et al., 2024; Zhang et al., 2023). Each M_0_, label, and control acquisition consisted of 12 spiral interleaves acquired over 12 TRs. Three dynamic image volumes were reconstructed for each acquisition. Additional imaging parameters included: acceleration rate (R) = 4, TR = 7 s, effective perfusion-weighted image (PWI) volume TR (TRPWI) = 56 s, echo spacing = 19.5 ms, TE = 1.72 ms, ADC bandwidth = 400 kHz, ADC duration = 10.5 ms, flip angle = 15°, and echo train length (ETL) = 72. We employed a novel reconstruction framework to image the brainstem by integrating compressed sensing reconstruction with motion-resolved self-navigation to reduce motion artifacts (Jung et al., 2009; Lustig et al., 2007). The inverse problem was solved using the alternating direction method of multipliers (ADMM) (Boyd, 2010), implemented with 10 ADMM iterations and three nested conjugate gradient steps per iteration. To correct for artifacts inherent to spiral acquisition, blurring was addressed using multi-frequency interpolation (MFI) (Man et al., 1997), and spiral trajectory errors were corrected by incorporating a gradient impulse response function (GIRF) measured on the scanner (Duyn et al., 1998; Robison et al., 2019; Vannesjo et al., 2013). To address the problem of B_0_ field inhomogeneity at 7T, we first conducted an ASL prescan (2:20 minutes) followed by B_0_ mapping scans (1:30 minutes) and the three ASL condition runs. The acquisition time of the pre-handgrip run was 10:16 minutes and for the subsequent handgrip and post-handgrip runs were 8:52 minutes.

### MRI data preprocessing

All anatomic and functional MRI data were automatically preprocessed using *fMRIprep* version 24.0.0 (Esteban et al., 2019) which is based on *Nipype* 1.8.6 (Gorgolewski et al., 2011). The complete *fMRIprep* boilerplate output is provided in the Supplementary Information. Subsequent preprocessing steps and statistical analyses were conducted using the AFNI software package (Cox, 1996). The two MT structural scans were analyzed using Advanced Normalization Tools, version 2.3.3 (Avants et al., 2009). Structural scans were aligned to the initial T1-weighted MP2RAGE scan using *antsRegistration*. A whole-brain group template was then generated through four iterations of *antsMultivariateTemplateConstruction,* including *N4BiasFieldCorrection*. All transformation matrices were concatenated and applied to subject-level MT structural scans via *antsApplyTransforms* to bring them from native space to group template space.

The eight functional scans, normalized to 2 mm isotropic MNI 152 non-linear 2009c asymmetric space via *fMRIprep*, were further smoothed with a 2 mm full-width half-maximum (FWHM) Gaussian kernel using AFNI’s *3dmerge* and scaled to 100% using AFNI’s *3dTstat* and *3dcalc*. A general linear model (GLM) was then estimated with AFNI’s *3dDeconvolve* to quantify condition-specific brain responses. Volumes exceeding 0.3 mm framewise displacement were censored to minimize motion artifacts (Ciric et al., 2017). The GLM included four stimulus conditions: standard, target, positive valence, and negative valence stimuli. To account for motion and physiological noise, the model incorporated six motion parameters, global signal regressors from white matter, cerebrospinal fluid, and the whole-brain mask, along with their temporal derivatives and quadratic terms resulting in a total of 36 nuisance regressors (Satterthwaite et al., 2013). The hemodynamic response function was modeled using a SPMG3 basis function which incorporates both the canonical response and its temporal and dispersion derivatives (Calhoun et al., 2004). To correct for temporal autocorrelation in the residuals and improve parameter estimation, we applied AFNI’s *3dREMLfit* using a voxelwise autoregressive-moving-average (ARMA) model proving more accurate 𝛽-parameter estimates by accounting for serial correlations in the fMRI time series.

ASL MRI data preprocessing was also performed using AFNI and ANTs. Linear motion correction of control and label images were conducted using AFNI’s *3dvolreg* and followed by registration of the mean control image to the T1-weighted anatomical image using AFNI’s *align_epi_anat.py*. A 12-parameter affine transformation was estimated from the anatomical image to the MNI152 template (mni_icbm152_nlin_asym_09c), and a subsequent nonlinear transformation was computed using ANTs *antsRegistration*.

### MRI data analyses

For functional MRI analyses, a LC consensus mask was applied to extract LC brain responses and cerebral blood flow (Dahl et al., 2022). We also utilized the publicly-available pontine reference area mask as a control region of interest (Dahl et al., 2022). Spatial smoothness was estimated using the autocorrelation function (ACF) of the residuals and averaged across subjects and sessions. Cluster thresholds for family-wise error correction (𝛼 = .05) were then determined using AFNI’s *3dClustSim* based on the averaged ACF parameters and were applied to statistical maps thresholded at a voxelwise criterion of *p* < .001. This approach accounts for the non-Gaussian nature of spatial autocorrelation in functional MRI data and follows the standards and threshold of best practices for cluster-level inferences (Cox et al., 2017). Using a nearest-neighbor connectivity of two and bi-sided voxelwise threshold of *p* = .001, simulations indicated that a minimum cluster extent of five contiguous voxels was required to achieve a family-wise error rate of *p* < .05. Only clusters meeting this threshold are reported. To observe activity and functional connectivity within the major attention networks in the brain, we used a seven-network parcellation previously delineated using large-scale intrinsic connectivity networks (Yeo et al., 2011). We used the strict mask version of the parcellation to minimize overlap and ensure precise assignment of voxels to the following networks: visual, dorsal attention, ventral attention, and frontoparietal attention networks. Finally, amygdala masks were derived from the Jülich Histological Atlas with a probability threshold of > 60% to ensure high anatomical specificity for each subregion. 𝛽-parameter estimates were extracted from all masks across all stimulus types using AFNI’s *3dmaskave*. When conducting post-hoc *t-* test analyses, we report the Holm-Bonferroni corrected *p-*values (Holm, 1979). In addition, if Mauchly’s test of sphericity was violated, we applied the Greenhouse-Geisser correction (Greenhouse & Geisser, 1959). Effect sizes for analysis of variance (ANOVA) analyses are reported as generalized eta squared (*η^2^*).

For structural MRI contrast analyses, we created a study-specific LC group mask to quantify intensity changes using the high resolution of our scans. First, MT structural scans were aligned to the T1-weighted MP2RAGE structural scans, and a group template was created using *antsMultivariateTemplateConstruction.sh*. The LC was then manually delineated in ITK-SNAP version 4.2 (Yushkevich et al., 2006) based on hyperintensity patterns that were reliably visible at 1 mm resolution (see Figure 8A). This group-level LC mask was subsequently transformed back into each participant’s native MT space using ANTs *applyTransforms* with the corresponding inverse warps. Voxelwise intensity values were extracted from each native-space LC mask using AFNI’s *3dmaskdrump*. LC voxels were normalized by dividing each voxel’s value by the mean signal intensity of a pons reference region restricted to the same z-slices as the LC mask. Finally, we conducted a clusterwise permutation analysis to test for significant differences in LC intensity between control and handgrip blocks, averaged across sessions (voxelwise threshold, *p < .*005; cluster-level family-wise error (FWE) correction, *p < .*05; 10,000 iterations).

In the ASL scans, to evaluate differences in cerebral blood flow between conditions, we used a nonparametric bootstrap randomization procedure with 10,000 iterations and 95% bias-corrected accelerated (BCa) confidence intervals (𝛼 = .05).

### fMRI Functional Connectivity Analyses

To examine dynamic interactions between the LC and the whole brain, we applied the 𝛽-series correlation approach (Rissman et al., 2004). For each subject and session, trial-wise GLM estimates were obtained using least-squares-all estimation with individual trial modulators (Mumford et al., 2012) from the residual maximum likelihood (REML) fit. Per-trial 𝛽-maps were generated for the LC seed as well as for the central pontine reference mask. Seed-to-voxel 𝛽-series correlations were computed separately for the control and handgrip conditions, with whole-brain Pearson correlations estimated using AFNI’s *3dTcorr1D.* Thus, each beta series reflects a voxel’s estimated activity during the presentation of each stimulus for each task condition. Correlation coefficients were converted to Z-scores via Fisher’s *r*-to-*z* transformation. Condition differences were computed using AFNI’s *3dttest++* with family-wise error correction at *p < .*05 and a voxelwise threshold of *p* < .005.

### Brain-behavior partial least square correlations (PLSC)

To investigate the relationship between brain activity and behavioral performance across sessions, we employed partial least squares correlation analyses, a multivariate statistical framework designed to identify patterns of covariance between two variables (Krishnan et al., 2011; McIntosh & Lobaugh, 2004), as previously used in LC neuroimaging research (Dahl et al., 2022). Unlike univariate approaches that test each association independently, PLSC examines the joint relationship across all variables simultaneously, making it robust for datasets across modalities. In this study, we implemented a cross-linked PLSC approach to integrate information from both scanning sessions into a single model rather than analyzing sessions independently. In this framework, data from both sessions were concatenated in a block structure and latent variables (LVs) were extracted to reflect patterns of shared variance consistent across sessions. This method enhances sensitivity to stable brain-behavior relationship by pooling variance across sessions, while also allowing for post-hoc comparisons to evaluate the magnitude and direction of these patterns per session. PLSC computes the cross-covariance matrix between measures and decomposes it via singular value decomposition (SVD) to identify the latent variables that maximize shared variance. Each LV includes a pair of weight vectors for each dataset and describes the contribution of each variable to the multivariate pattern. In this study, we explored the relationship between LC brain responses to each stimulus type (standard, target, positive valence, negative valence) for each hemisphere with recognition memory hit rates for emotionally valenced faces during the recognition memory task. To evaluate statistical robustness, we performed permutation testing (10,000 iterations) to assess the significance of each LV and bootstrap resampling (10,000) iterations to estimate the stability of individual variable weights. Bootstrap ratios (BSRs) were calculated as the ratio of each weight to its bootstrap-derived standard error and were interpreted analogous to Z-scores, in which |BSR| > 1.96 was deemed statistically significant.

### Data Availability Statement

The datasets generated during the current study are available in the *OpenNeuro* repository (ds006690; repository will be made public after acceptance).

## Results

### Arousal induced by handgrip increases cerebral blood flow to the LC

We first examined whether the physiological arousal manipulation induced by isometric handgrip selectively modulated LC metabolic activity in the ASL pulse sequences. To contextualize these effects, we included the pons as a neighboring reference region for cerebral blood flow (CBF) and the amygdala to examine the specificity of the arousal manipulation. Arousal significantly increased CBF in the LC during the handgrip period relative to the pre-handgrip period, 95% CI [2.76, 11.62], *p <* .001 (pre-handgrip: mean = 37.8, SE = 3.18, handgrip: mean = 43.8, SE = 3.32; see Figure 2), but not in the adjacent pontine reference region, 95% CI [-3.38, 7.02], *p =* .642 (pre-handgrip: mean = 34.6, SE = 3.59, handgrip: mean = 36.0, SE = 3.22). In contrast, physiological arousal significantly reduced CBF in the amygdala, 95% CI [-4.47, -1.67], *p <* .001 (pre-handgrip: mean = 33.9, SE = 0.94; handgrip: mean = 30.6, SE = 0.54). In addition, CBF did not differ during the handgrip and post-handgrip period in the LC, 95% CI [-12.53, 0.69], *p =* .061 (post-handgrip: mean = 37.4, SE = 4.55), pontine reference region, 95% CI [-3.65, 1.00], *p =* .212 (mean = 34.5, SE = 2.73), and amygdala, 95% CI [-1.00, 2.21], *p =* .331 (mean = 31.4, SE = 0.80). Quantitative cerebral blood flow maps are shown in the Supplementary Information.

**Figure 2.**
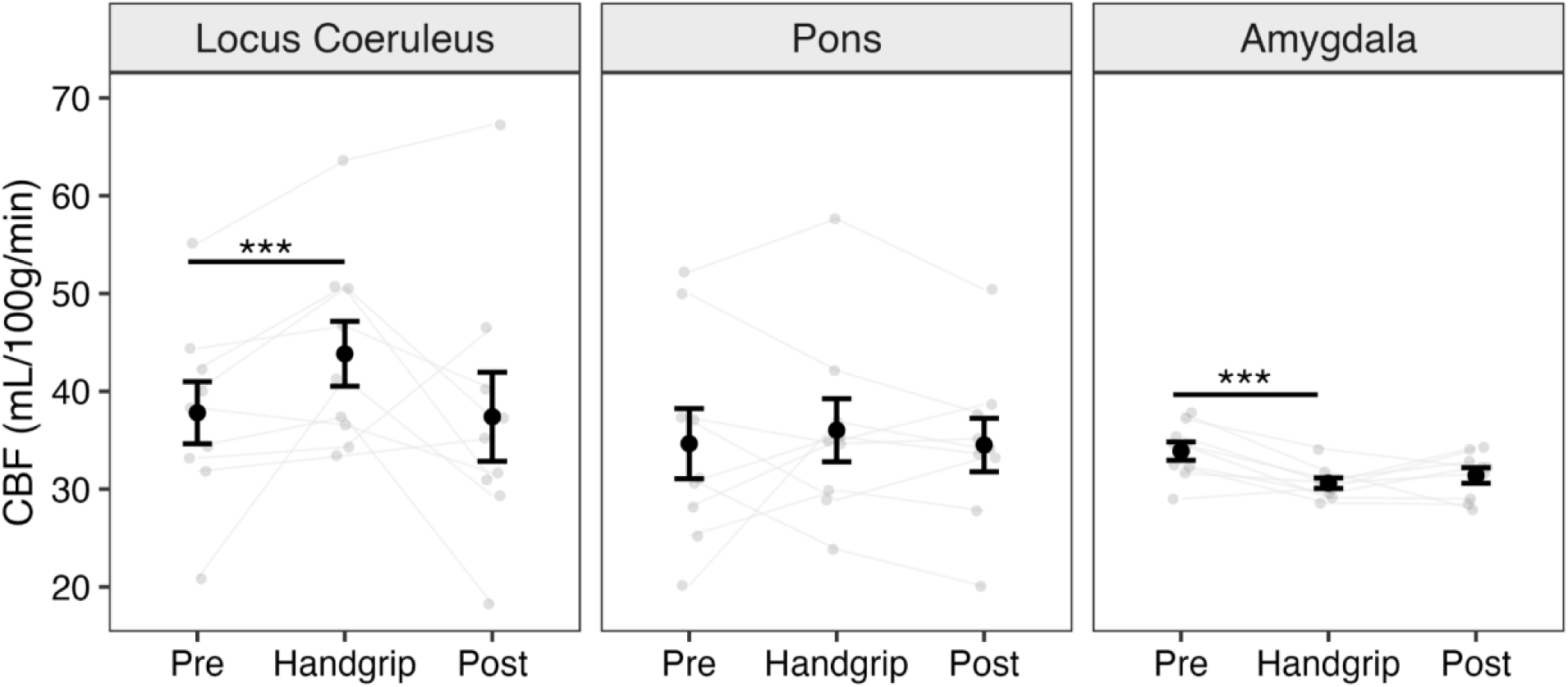
Arousal specifically increases metabolic activity in the LC. In nine participants, we examined how handgrip modulated metabolic activity and whether it persisted in the post-handgrip rest period using a nonparametric bootstrap randomization procedure with 10,000 iterations. Arousal increased cerebral blood flow (CBF) to the LC, *p <* .001, but not in the pons. In addition, arousal decreased CBF to the amygdala, *p <* .001. Finally, we identified no differences in the handgrip and post-handgrip period for the LC, pons, and amygdala.

### Arousal increases LC blood oxygen level-dependent (BOLD) responses to target and angry faces, but not happy faces

Given that handgrip specifically increased metabolic activity in the LC, we examined whether these arousal-related changes also modulated functional LC responses. A repeated-measures analysis of variance (RM ANOVA) analysis with factors condition (control, arousal), session (1,2), and stimulus (standard, target, positive, negative) revealed significant main effects of condition, *F*(1,21) = 5.82, *p* = .025, *η^2^_g_* = .045, and stimulus, *F*(2.75,57.84) = 19.74, *p* < .001, *η^2^g* = .063, but not of session, *F*(1,21) = .75, *p* = .397. Critically, a condition x stimulus interaction was significant, *F*(3,63) = 4.80, *p* = .004, *η^2^_g_* = .015 (see Figure 3A), indicating that arousal differentially modulated LC responses depending on stimulus type. Post-hoc *t*-test comparisons, corrected for multiple comparisons, indicated that LC BOLD responses were greater in the arousal condition compared with the control condition for target stimuli, *t*(21) = 3.40, *p* = .003, *d* = .741, and negative stimuli, *t*(21) = 2.59, *p* = .017, *d* = .565, but not for standard or positive stimuli, *ts*(21) < 1.27, *ps* > .218, supporting our hypothesis. Within the control condition, responses to target stimuli exceeded those to standard stimuli, *t*(21) = 2.88, *p* = .041, *d* = .628, but not to any other stimuli, *ts*(21) < 2.59, *ps* > .074. In the arousal condition, responses to the standard stimuli were reduced relative to target, *t*(21) = 9.74, *p* < .001, *d* = 2.12, positive, *t*(21) = 5.26, *p* < .001, *d* = 1.147, and negative stimuli, *t*(21) = 6.50, *p* < .001, *d* = 1.419. Finally, target stimuli evoked greater responses than positive stimuli in the arousal condition, *t*(21) = 3.69, *p* = .007, *d* = .805. See the Supplementary Information for visualizations of the BOLD response within the brainstem. In addition, we did not identify any main effects of condition or interactions in any of the subdivisions of the amygdala (also see Supplementary Information).

**Figure 3.**
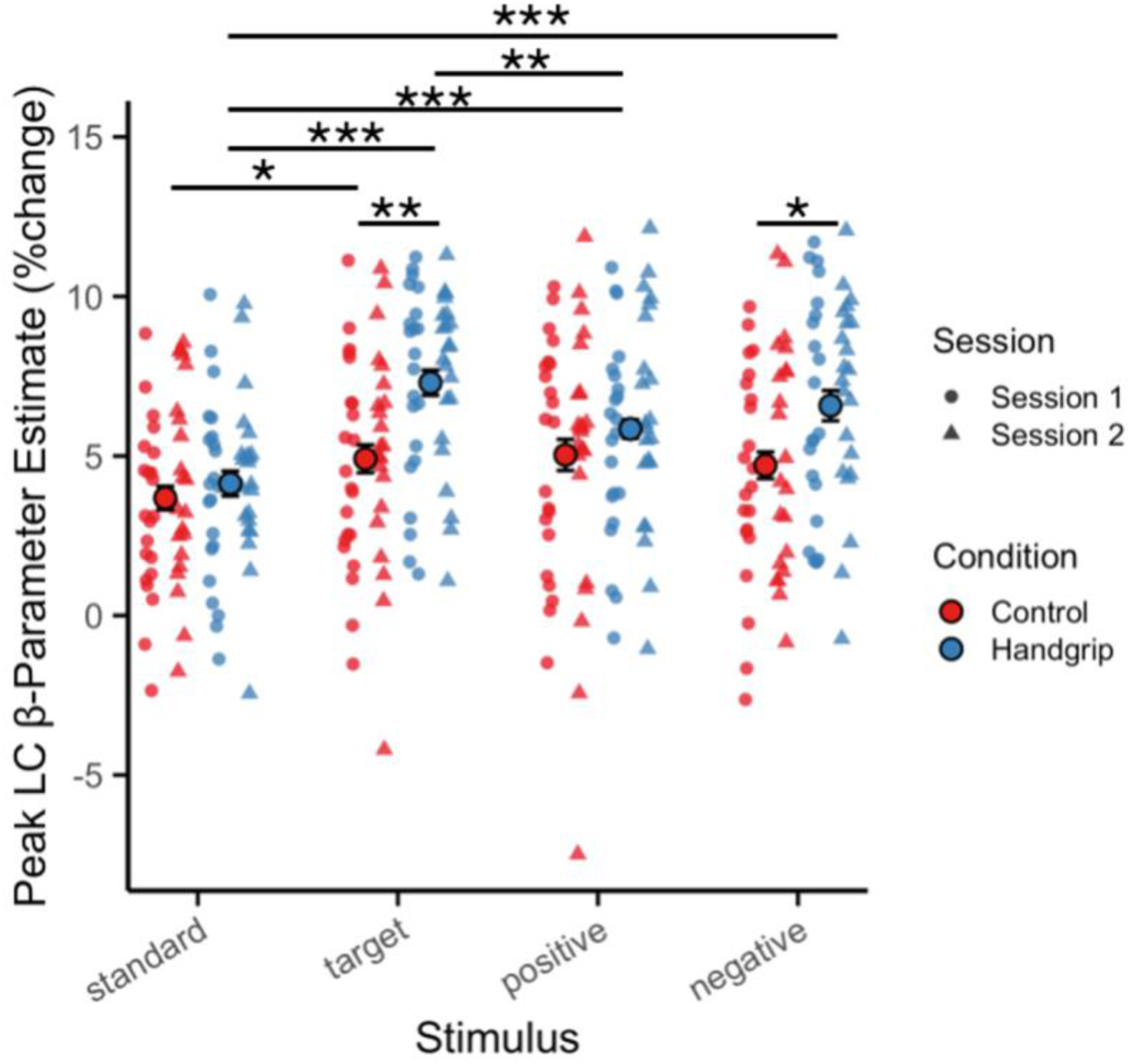
Arousal increases brain responses to prioritized and negative-valence stimuli in the LC. Given the small number of voxels encompassed by the LC mask at 2 mm resolution (left = 6, right = 4) and the susceptibility to noise from a single misaligned voxel (Yi et al., 2023), we extracted the peak voxel within the LC for 𝛽-parameter estimates across two sessions. A repeated-measures ANOVA revealed a significant condition x stimulus interaction that was further explored using *t*-test comparisons. During the handgrip block, LC responses were further elevated when processing target and angry faces but not for happy faces. Full descriptive statistics are reported in the Supplementary Information.

### Reduced LC and default mode network connectivity during target and angry face processing

Given that LC activity modulates large-scale cortical networks (Dahl et al., 2025; Shine et al., 2016; van den Brink et al., 2019), we examined network interactions between the brainstem and the whole brain using a trial-level 𝛽*-*series correlation analysis (Rissman et al., 2004), with the LC as a seed region across all stimulus types. Within the control condition, we contrasted connectivity maps for standard – target, standard – positive, and standard – negative faces to evaluate differences in connectivity associated with prioritized and emotional stimuli. When processing target and angry faces, significant clusters within nodes of the default mode network were suppressed in the target – standard contrast (peak Z = -3.28, left vmPFC, 6 voxels, [6.5, -61.5, -4.5]) and the negative – standard contrast (peak Z = -3.09, right vmPFC, 5 voxels, [-11.5, -47.5, 1.5]; peak Z = -3.37, left dorsal striatum, 5 voxels, [26.5, -7.5, 13.5]; peak Z = -3.16, right posterior angular gyrus, 5 voxels, [-53.5, 70.5, 29.5]). However, no significant clusters were identified for the positive – standard contrast, underscoring a valence-dependent pattern of LC functional connectivity (see Figure 4). To check whether these functional connectivity results were specific to the LC, we conducted the identical analyses with the central pontine reference region as the seed, and no significant clusters were identified in the FWE-corrected contrasts.

**Figure 4.**
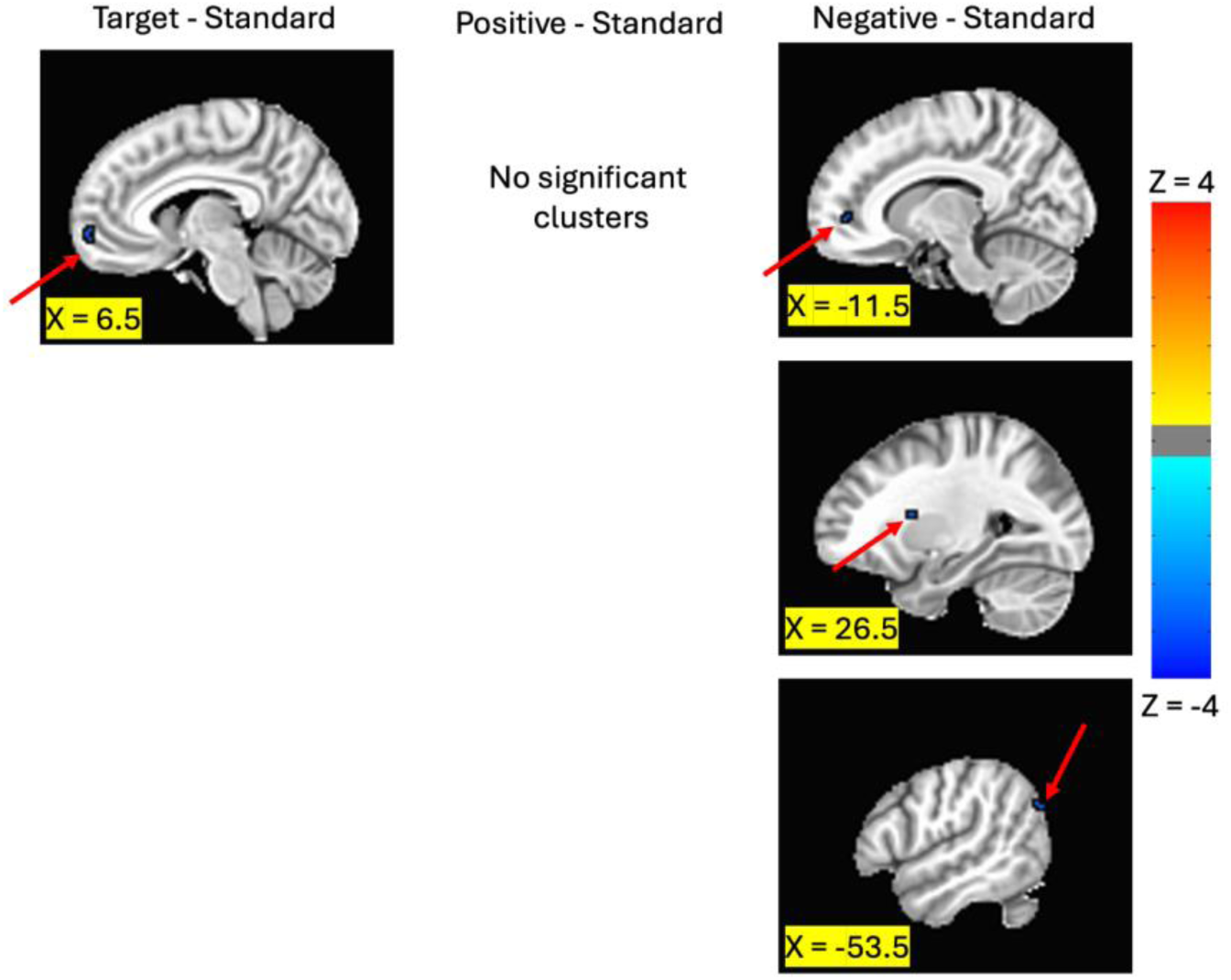
Valence dependent decoupling of LC activity from the default mode network. We conducted trial-level 𝛽*-*series correlation analysis in twenty-two participants, using the LC as a seed region to examine connectivity with the whole brain. Compared with standard stimulus processing, target and negative-valence stimuli were associated with reduced connectivity between the LC and nodes of the default mode network, including the ventromedial prefrontal cortex, left dorsal striatum, and right posterior angular gyrus.

### Arousal differentially modulates stimulus responses across attention networks

Because arousal selectively modulated 𝛽-parameter estimates in the LC but not in amygdala subdivisions, we next examined whether arousal differentially influences activity within attention networks that are modulated by the LC (Corbetta et al., 2008). Using intrinsic functional connectivity maps defined by Yeo et al. (2011), we tested the effects of arousal within functional network parcellations of the visual system, dorsal attention network (DAN), ventral attention network (VAN), and frontoparietal network (FPN). A RM ANOVA with condition (control, arousal), session (1,2), stimulus (standard, target, positive, negative) and region (visual, DAN, VAN, FPN) as factors revealed significant main effects of stimulus, *F*(2.01,42.20) = 19.51, *p* < .001, *η^2^g* = .018, and region, *F*(2.54,53.30) = 3.45, *p* = .029, *η^2^g* = .029, but not for condition, *F*(1,21) = 2.90, *p* = .103, and session, *F*(1,21) = .97, *p* = .337. In addition, the three-way condition x stimulus x region interaction, *F*(5.00,105.09) = 2.66, *p* = .026, *η^2^_g_* = .004, and the three-way session x stimulus x region interaction, *F*(5.99,125.89) = 2.67, *p* = .018, *η^2^_g_* = .003, were significant (see Figure 5). To further examine these interactions, we conducted separate RM ANOVAs with condition, session, and stimulus as factors in each region.

**Figure 5.**
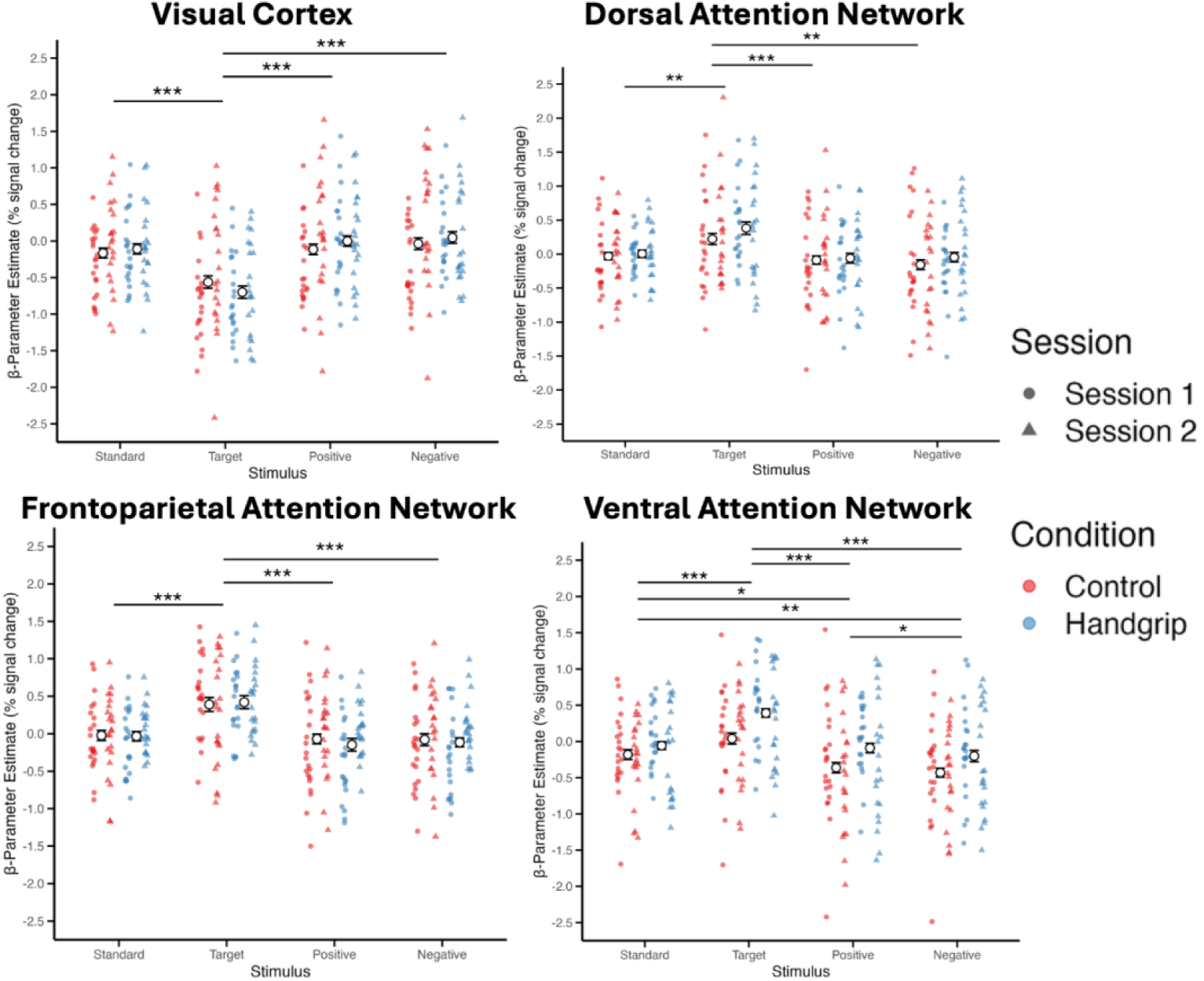
Arousal enhances salience network responses and differentially modulates visual cortex activity by stimulus type. Mean parameter estimates were computed across all intrinsic functional connectivity maps defined by Yeo et al. (2011). In the visual system, arousal further suppressed target responses while increasing responses to all other stimuli. We identified no effects of arousal in the dorsal attention and the frontoparietal attention networks. In the ventral attention network, arousal increased responses to all stimuli. Means and standard errors reflect the average across both sessions. *\*p* < .05, *\*\*p* < .01, *\*\*\*p < .*001.

In the visual network, we identified significant main effects of session, *F*(1,21) = 5.78, *p* = .025, *η^2^_g_* = .034, stimulus, *F*(2.17, 45.55) = 76.23, *p* < .001, *η^2^_g_* = .139, but no main effect of condition, *F*(1,21) = .06, *p* = .810. Target stimuli evoked significantly reduced responses compared with all other stimuli, *ts*(21) > 8.68, *ps* < .001, *ds* > 1.850. In addition, a significant condition x stimulus interaction, *F*(2.88,60.54) = 3.85, *p* = .015, *η^2^_g_* = .006, indicated that arousal further suppressed target responses while increasing responses to all other stimuli. In the dorsal attention network, we observed a significant main effect of stimulus, *F*(1.64, 34.41) = 14.76, *p* < .001, *η^2^_g_* = .069, but no main effects of condition, *F*(1,21) = .83, *p* = .373, or session, *F*(1,21) = .14, *p* = .708. Post-hoc analyses showed that target stimuli elicited greater responses than all other stimuli, *ts*(21) > 3.57, *ps* < .007. *ds* > .761. In the ventral attention network, we identified a significant main effect of condition, *F*(1,21) = 7.59, *p* = .012, *η^2^_g_* = .035, reflecting higher activity in the arousal than in the control condition. There also was a significant main effect of stimulus, *F*(3,63) = 35.61, *p* < .001, *η^2^_g_*= .090, but not of session, *F*(1,21) = 1.28, *p* = .271. Post-hoc analyses revealed that target stimuli evoked greater responses than all other stimuli, *ts*(21) > 5.31, *ps* < .001, *ds* > 1.132, and both emotionally-valenced stimuli showed suppressed responses compared with standard stimuli, *ts*(21) > 2.43, *ps <* .024, *ds* > .519. In the frontoparietal attention network, we identified a significant main effect of stimulus, *F*(1.78, 37.44) = 37.16, *p* < .001, *η^2^_g_* = .146, but no main effect of condition, *F*(1,21) = .04, *p* = .836, nor session, *F*(1,21) = 2.44, *p* = .134. Post-hoc analyses revealed that target stimuli had significantly greater responses compared with all other stimuli, *ts*(21) > 6.42, *ps* < .001, *ds* > 1.368. In addition, there was a significant session x stimulus interaction, *F*(2.68,56.23) = 12.88, *p* < .001, *η^2^_g_* = .021. In Session 1, the target stimuli produced significantly greater activation than all other stimuli, *ts*(21) > 8.01, *ps* < .001, *ds* > 1.75, and the positive and negative stimuli were also suppressed compared with the standard stimuli, *ts*(21) > 2.54, *ps* < .038, *ds* > .550. However, in Session 2, the target stimuli again stimulated greater activation than all other stimuli, *ts*(21) > 3.69, *ps* < .005, *ds* > .810, but the emotional stimuli were not suppressed compared with the standard stimuli, *ts(*21) < .22, *ps* = 1.000. Full descriptive statistics for all regions are reported in the Supplementary Information.

### Arousal during encoding increases recognition memory for angry, but not happy, faces

Following the oddball task portion of the experiment, participants completed a recognition memory task that included all previously presented emotional faces as well as novel lures to assess false positives. We examined whether memory for valenced faces was differentially modulated by the arousal condition. Based on prior findings that LC activity enhances memory under arousal (Clewett et al., 2018), we predicted that negative-valence faces presented during the arousal block would be better recalled. Sensitivity (*d*’) reflects the extent to which participants can discriminate between previously presented and novel items, with higher values indicating more accurate discrimination (Stanislaw & Todorov, 1999). We conducted a RM ANOVA with session (1,2), condition (control, arousal), and valence (positive, negative) as factors. Critically, the two sessions enabled us to capture distinct attentional processes engaged by novel emotional faces. In Session 1, participants were unaware that a recognition test would follow and suppressed task-irrelevant emotional stimuli while only counting target oddball faces. In Session 2, participants were aware of the recognition test and allocated attention to the emotional stimuli.

We identified a significant main effect of session, *F*(1,21) = 7.05, *p = .*015, *η^2^_g_* = .064, indicating an overall higher sensitivity in Session 2 compared with Session 1. In addition, no main effects were observed for condition, *F*(1,21) = 1.45, *p* = .242, or valence, *F*(1,21) = .24, *p* = .524. Critically, a three-way session x condition x valence interaction was significant, *F*(1,21) = 5.36, *p* = .031, *η^2^_g_*= .014 (see Figure 6). Post-hoc comparisons revealed no performance differences for prior faces presented in the control block (Session 1: *t*(21) = .20, *p* = .842; Session 2: *t*(21) = -.97, *p* = .345). However, for faces presented during the handgrip block, there were no valence effects in Session 1, *t*(21) = 1.69, *p* = .105, but in Session 2 when paying attention to the emotional faces, sensitivity toward negative-valence faces were significantly higher than to positive-valence faces, *t*(21) = -2.21, *p* = .038, *d* = .472.

**Figure 6.**
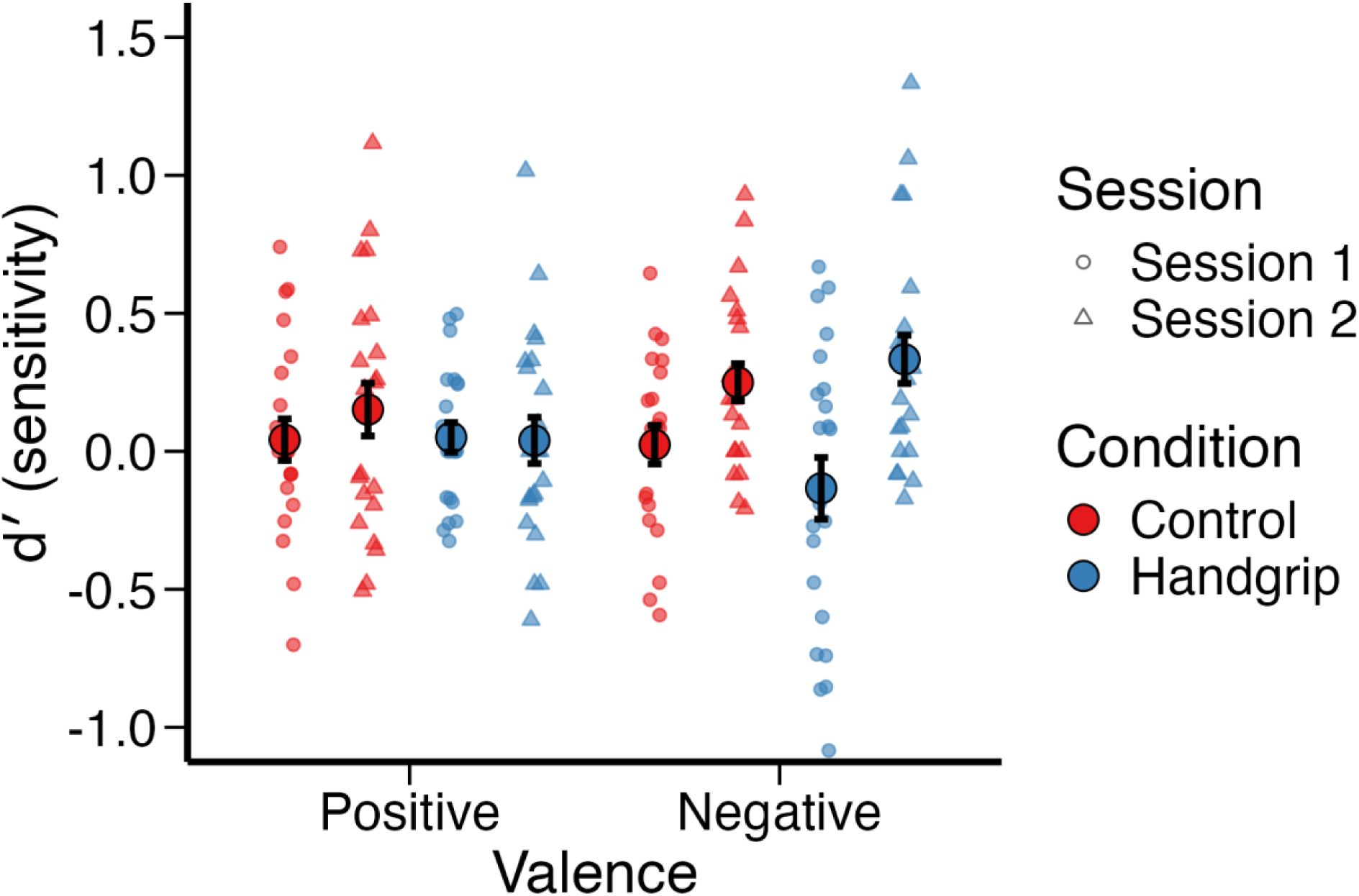
Arousal modulated valence-specific effects in recognition memory performance. In Session 1, participants were unaware of the upcoming recognition memory test and focused on counting target faces while suppressing task-irrelevant faces, resulting in no significant valence effects. However, in Session 2, participants were aware of the recognition memory task and allocated attention to the emotional faces. We conducted a repeated-measures ANOVA over factors session, valence, and condition with twenty-two participants and identified a significant three-way interaction for sensitivity. Memory accuracy (d‘) was significantly higher for angry relative to happy faces presented during the handgrip block in Session 2, but there were no significant differences observed for faces presented during the control block.

### Suppressing and attending to emotional faces is coupled with left LC BOLD responses

To examine multivariate relationships between LC responses and behavior, we implemented partial least squares correlation (PLSC) which uses singular value decomposition (SVD) to assess shared covariance between datasets (Dahl et al., 2022; Krishnan et al., 2011; McIntosh & Lobaugh, 2004). Specifically, this method identifies latent variables (LVs), or weighted combinations of the original variables, that optimally capture shared information between data modalities. Because our design included two sessions, we modeled session as a separate condition in the cross-block PLSC analysis, allowing us to capture session-specific covariance patterns rather than averaging across sessions. In this framework, each session generates its own LVs which are then stacked together and interpreted within a common model, thus allowing estimates of brain-behavior relationships separately for each session while also allowing for their joint interpretation. We entered recognition memory hits for emotional stimuli as the behavioral measure (X matrix) and LC brain responses to all stimuli as the brain measure (Y matrix).

The cross-block PLSC analyses revealed a single significant LV that accounted for 81.6% of the cross-block covariance, *p* = .020, (see Figure 7A). This LV optimally expressed the multivariate association between LC brain responses and behavioral performance, *r* = .470, *p* = .001 (see Figure 7B), demonstrating that the global covariance structure is represented by one positive direction. Interestingly, only left LC brain responses to target, and both emotional stimuli reliably contributed to this latent variable as defined by bootstrap ratio (BSR] cutoffs > |1.96| (see Figure 7C). We next stratified the PLSC model by session. Spearman correlations between the LV brain scores and behavioral performance showed opposite patterns across sessions (see Figure 7D). In Session 1, weaker left LC activation predicted higher recognition accuracy, while in Session 2, the same LC response pattern predicted higher accuracy. This context-dependent shift demonstrates that the coupling between LC activity and memory performance shifted as participants transitioned from suppressing task-irrelevant distractors to allocating more attention to emotional stimuli (as they now knew they would be tested on their memory for these).

**Figure 7.**
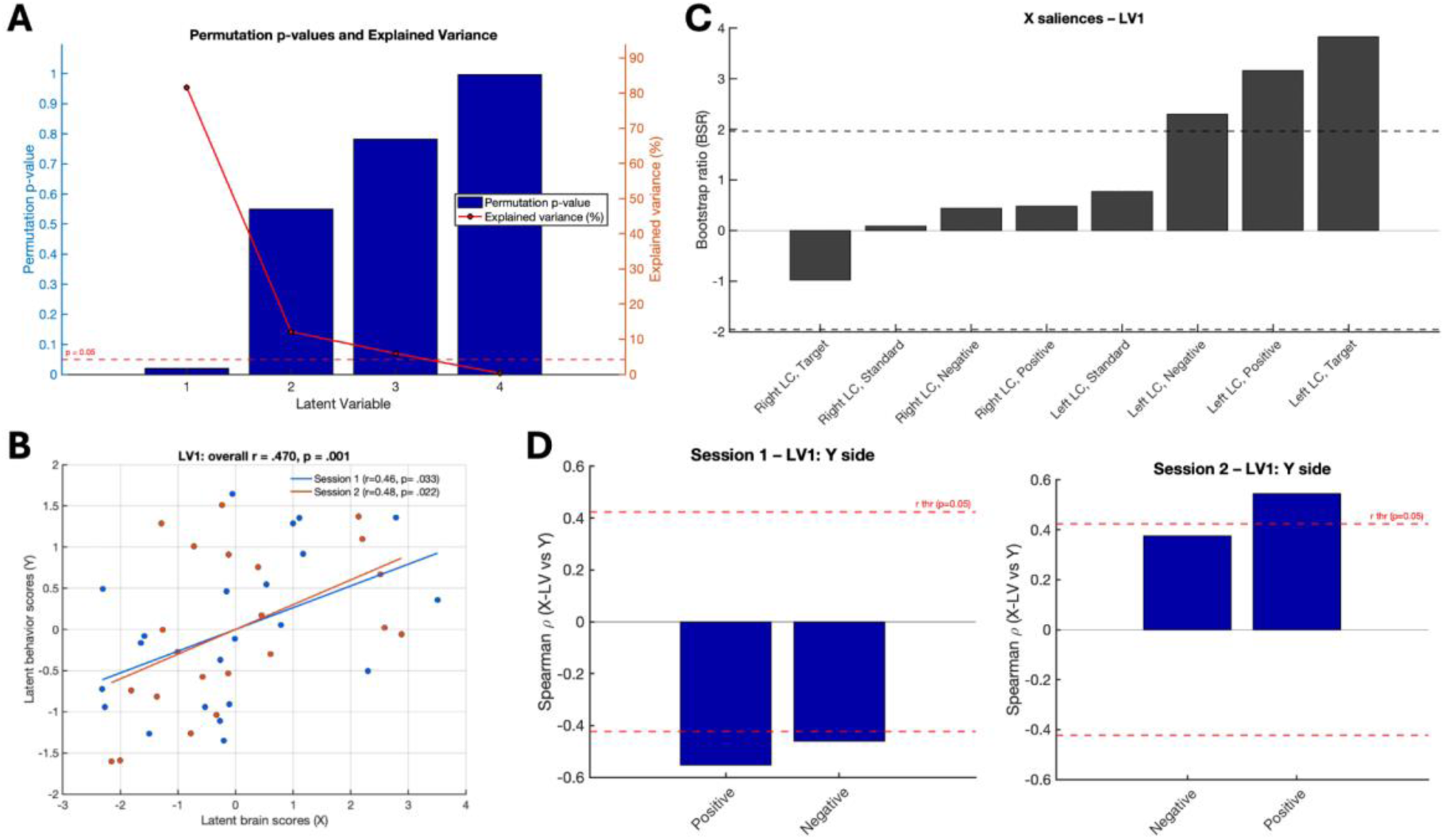
Left LC brain responses are linked to recognition memory performance. We conducted cross-block partial least squares correlation to examine brain-behavior relationships between the LC and recognition memory. (A) The analysis identified a single latent variable (LV) that explained 81.6% of the cross-block covariance. (B) Both latent behavioral and brain scores loaded significantly on this LV. (C) Bootstrap ratios indicated that only left LC brain responses to target, happy, and angry faces significantly contributed to this LV. Given that partial least square correlation modalities are sign-indeterminate, we inverted the bootstrap ratio and correlation values for better intuitive visualization, such that greater left LC brain responses mapped onto positive loadings. (D) Post-hoc Spearman correlations stratified by session revealed opposite brain-behavior associations. Correct recognition memory to emotional faces were linked to weaker left LC responses in Session 1, but stronger responses in Session 2. These results show that LC-behavior coupling is context dependent and reflect shifts between attention suppression and allocation.

### Arousal decreases LC MRI contrast in magnetization transfer (MT) structural scans

Finally, we investigated whether short-term changes in arousal modulated LC MRI contrast in MT structural scans. Currently, LC MRI contrast is a frequently used measure of LC integrity in humans (Berger et al., 2023; Betts et al., 2019; Bueichekú et al., 2024; Dahl et al., 2023, 2022). However, information regarding whether this structural imaging signal is affected by acute physiological changes is lacking, despite the fact that current models suggest that LC MRI contrast is primarily driven by signals from protons in fluid interacting with signals from macromolecules (Henkelman et al., 2001), properties that could be quickly altered depending on changes in fluid dynamics in a region. Thus, we examined whether LC MRI contrast is sensitive to transient physiological states. We predicted that increases in arousal would be accompanied by changes in LC MRI contrast. During the handgrip block, participants were required to continuously squeeze the handgrip throughout the duration of the scan. In both the left and right LC, handgrip was associated with decreased MRI contrast relative to control (see Figure 8B). In the left LC, three significant clusters were identified that were 8, 6, and 17 voxels with peak *t*-statistic of -4.308, -4.658, and -4.445, respectively (mask i,j,k locations [3.5,1.5,2.5], [4.5,0,3], [2,1,5]). In the right LC, two significant clusters of 5 and 12 voxels were identified with peak *t-* statistic of -4.705 and -6.349, respectively (mask i,j,k locations [2.5,1.5,5] and [3,1.5,7]). These condition differences could not be attributed to motion as quantified by MRIQC’s image quality metrics (see Supplementary Information; Esteban et al., 2017).

**Figure 8.**
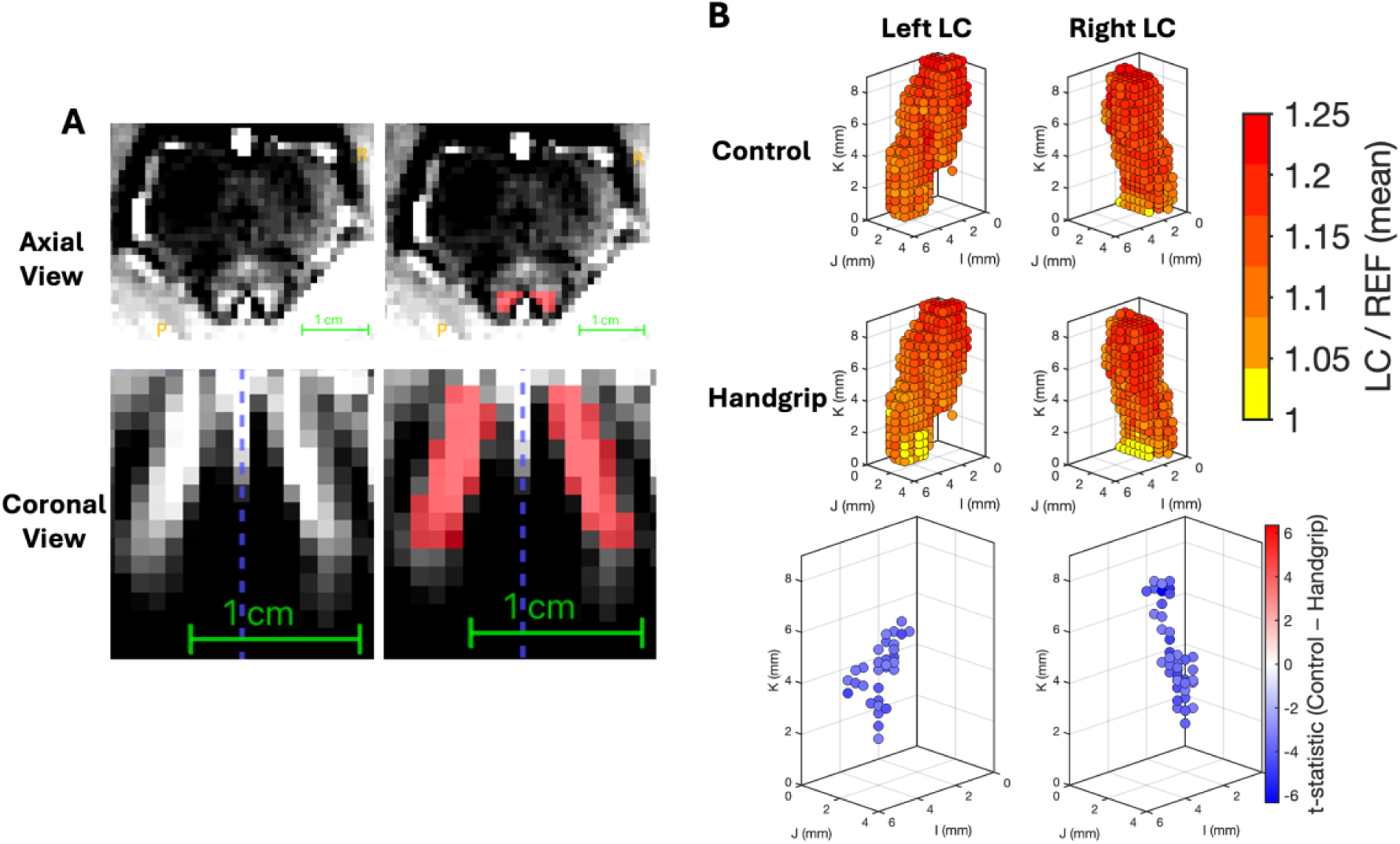
Handgrip decreases LC MRI contrast compared with control conditions. (A) A study-specific LC group mask was created by manually delineating hyperintensities in the group template derived from high-resolution MT structural scans. (B) For voxelwise cluster-based analyses, each LC voxel’s MRI contrast was normalized by the mean intensity of a central pontine reference region matched in the number of z-slices to the LC mask. Analyses revealed significantly reduced LC MRI contrast during handgrip scans compared with control scans. Significant clusters from the voxelwise permutation test are highlighted in the Control - Handgrip contrast.

## Discussion

In this study, we tested whether the LC serves as the primary driver of valence-specific effects within affective attention networks. Using an isometric handgrip paradigm to elicit arousal (Bachman et al., 2023; Mather et al., 2020; Nielsen & Mather, 2015), we observed selective increases in cerebral blood flow to the LC while adjacent brainstem regions such as the pons remained unaffected. In addition, the amygdala showed decreases in cerebral blood flow, underscoring the specificity of LC engagement in increasing metabolic activity. The specificity of our arousal manipulation to the LC reflects the direct innervation from autonomic brainstem nuclei, including the nucleus paragigantocellularis (nPGi) and the nucleus of the solitary tract (NTS) (Mello-Carpes & Izquierdo, 2013). Baroreceptors in the carotid sinus and aortic arch detect interoceptive blood pressure changes and transmit signals via cranial nerves to the NTS, which projects excitatory inputs to the LC (Dampney, 2015; Huang et al., 2025). Unlike the LC, the amygdala does not receive direct baroreceptor inputs but processes arousal regulations through its interactions with the LC and hypothalamus. Thus, we interpret the observed arousal effects as mediated by the LC through its widespread noradrenergic projections on cognition and attention.

By combining ultra-high field imaging with advanced reconstruction methods (Woodward et al., 2023), we demonstrated that cerebral blood flow provides a noninvasive metabolic index of LC function. We implemented three methodological advances to develop a novel, optimized whole-brain perfusion approach that includes the brainstem at 7T: (1) a pseudo-continuous ASL (pCASL) sequence with comprehensive optimizations for 7T MRI (Wang et al., 2022); (2) a Fast Low-Angle Shot (FLASH)-based readout with rotated golden-angle stack-of-spirals (rGA-SoS) sampling (Guo et al., 2024; Munsch et al., 2020; Zhao et al., 2023); and (3) dynamic compressed sensing (CS) reconstruction that enables high spatiotemporal resolution and incorporates self-navigation for motion correction (Jung et al., 2009; Lustig et al., 2007). Typical BOLD MRI scans do not provide quantitative metrics of brain activity (Gore, 2003). Activity in response to a stimulus or event is contrasted with events at other times during the scan sequence (even this has issues due to signal drift; e.g., Yan et al., 2009), but the absolute value of the signal at any particular point in time cannot meaningfully be compared across participants or scans. In contrast, ASL provides a quantitative estimate of cerebral blood flow values that can be compared not only across conditions within a scan but also across scans and across people producing absolute measures of perfusion (Pike, 2012). However, with 3T MRI, the spatial resolution of ASL sequences does not allow for assessing signal from brainstem nuclei (Clement et al., 2022). Our novel 7T pCASL sequence establishes the first non-invasive neuroimaging approach to assess the absolute level of LC activity in humans. Thus, these high-resolution cerebral blood flow measures should provide an important addition to current fMRI, pupillometry and electrophysiology approaches to assess LC function (Costa & Rudebeck, 2016; Joshi & Gold, 2020; Nieuwenhuis et al., 2005; Weiss et al., 2025).

In our functional data, we observed that the LC responded more strongly to prioritized stimuli than to standard stimuli in the emotional oddball task under control conditions. When we induced arousal through handgrip, LC brain responses further increased to target stimuli with high attentional priority and to negative-valence stimuli. Consistent with our cerebral blood flow findings, arousal did not enhance amygdala responses in any subdivision. Importantly, both angry and happy faces elicited greater amygdala activity than target or standard stimuli across both control and handgrip conditions, replicating the valence-general conclusions in the literature (Costafreda et al., 2008; Lin et al., 2020; Sergerie et al., 2008). Functional connectivity analyses revealed that target and negative stimuli, relative to standard stimuli, decoupled LC activity from the default mode network (DMN), unlike with positive stimuli. Prior work has shown that the DMN deactivates during engagement with salient, goal-directed or emotional stimuli (Dahl et al., 2025; Raichle et al., 2001), and often exhibits anti-correlated activity with arousal and attention networks (Kucyi et al., 2017; Sridharan et al., 2008). In support of this, we found that arousal significantly modulated activity in attention systems. The salience network showed a general increase in activity across all stimuli in the handgrip block, whereas the visual system demonstrated an interaction in which arousal further suppressed responses to target stimuli but enhanced responses to other stimulus categories. Behaviorally, participants recognized negative faces more accurately than positive faces under arousal and multivariate analyses linked recognition performance to left LC brain activity. Taken together, these findings indicate that LC arousal modulation is associated with increased salience network responses and reduced coupling with the DMN, which relates to biased attentional priority toward prioritized and negative-valence information.

An important aspect of our experiment design is that participants did not squeeze the handgrip during the functional scans. We aimed to avoid confounding the results with motor activity or cognitive load and instead leveraged the short-term physiological, residual effects of sustained handgrip exertion performed during the structural MT scan. Our ASL data confirmed that LC cerebral blood flow did not return immediately to baseline after handgrip but remained somewhat elevated during the approximately nine minute resting period, consistent with a sustained arousal response. Prior evidence similarly shows that muscle sympathetic nerve activity remains elevated during the post-handgrip ischemia phase before gradually returning to baseline during recovery (Boulton et al., 2019). Accordingly, our functional data reflect carryover effects of arousal on attention biases rather than concurrent increases in task demands. To control for these effects and isolate the arousal manipulation, we fully counterbalanced the sequence of control and handgrip blocks across participants and sessions. Finally, we interpret these effects as reflecting short-term potentiation and noradrenaline release rather than consequences of noradrenergic depletion, which typically emerges under prolonged stressors or administration of pharmacological blockers (Prinzi et al., 2023).

Structural LC imaging has emerged as one of the most widely used approaches in the fields of aging and neurodegenerative disease. Prior cross-sectional age-comparative findings show that LC MRI contrast increases across adulthood, peaks in midlife, and subsequently declines in older age (Liu et al., 2019; Riley et al., 2023). Furthermore, reductions in LC MRI contrast have been linked to neurodegenerative conditions such as Alzheimer’s disease and Parkinson’s disease (Betts et al., 2019; Liu et al., 2025; Madelung et al., 2022). One study has also shown that higher LC MRI contrast predicts stronger physiological arousal responses to stress (Bachman et al., 2022). In contrast, we observed that isometric handgrip decreased LC MRI contrast. Two factors are important for interpreting this result. First, rather than relying on a single voxel reflecting peak LC contrast that is typical in the literature, we used a cluster-based nonparametric permutation approach to identify significant clusters of change between blocks. Second, it is critical to consider what LC MRI contrast biologically reflects. Initial studies have attributed this structural signal to neuromelanin deposition in LC neurons, with age-related increase interpreted as accumulation of neuromelanin (Betts et al., 2019; Liu et al., 2019; Shibata et al., 2006). However, recent work by Watanabe and colleagues demonstrated that MT-weighted LC signal reflects a combination of neuronal properties including the high intracellular water content, relatively low macromolecular density, and the abundance of paramagnetic ions in catecholaminergic neurons, rather than T_1_ shortening molecules such as neuromelanin (Watanabe, 2023; Watanabe, Tan, et al., 2019; Watanabe, Wang, et al., 2019). These neuronal properties make the LC appear relatively bright on MT-weighted imaging because its signal is less suppressed than that of surrounding brainstem tissue. Moreover, this MT signal is also detectable in rodent LC despite the absence of neuromelanin (Watanabe, Wang, et al., 2019). In our study, handgrip increased cerebral blood flow to the LC, and perfusion imaging is sensitive to changes in the free-water proton pool associated with blood delivery and exchange (Le Bihan, 2007). Although the high intracellular water content in LC neurons leads to the elevated MT-weighted signal, blood contains mostly free water with very few macromolecules. Consequently, when blood flow increases to the LC, the added contribution of blood signal mixes with tissue signal, reducing the LC’s relative brightness by diluting its contrast against surrounding tissue. Therefore, we interpret our observed reductions in LC MT signal not as evidence of structural degeneration, but as reflecting an acute increase in local water content driven by elevated blood flow during short-term handgrip administration.

While amygdala hyperactivity is the most consistent finding across affective disorders (Etkin & Wager, 2007; Ressler & Mayberg, 2007), our findings point to a more prominent role of the neuromodulatory LC-NA system in shaping valence-specific attention biases. Dysregulation of the LC-NA system has been observed in anxiety disorders (Morris et al., 2020) and post-traumatic stress disorder (Boukezzi et al., 2025; Southwick et al., 1999). Post-mortem studies also identified that individuals with major depressive disorder exhibit elevated corticotropin-releasing factor in the LC (Bissette et al., 2003), indicating a chronic stress-related hypervigilant state of tonic, or baseline, neuromodulatory activity. These findings indicate that alterations in LC function contribute to psychiatric disorders. In rodents, optogenetic and chemogenetic stimulation of LC fibers modulated basolateral amygdala neuronal activity which induced anxiety-like behavior (Bayer et al., 2025; McCall et al., 2017), showing a mechanistic connection in which the amygdala functions as a key downstream target through which LC hyperactivity exerts pathological influence. Taken together, these findings highlight the LC as a critical yet underappreciated structure in affective disorders and underscores the critical need to clarify the mechanisms through which it contributes to models of psychopathology.

Our findings provide converging evidence that the locus coeruleus, and not the amygdala, serves as the key driver of arousal-driven, valence-specific effects within affective attention networks. By combining ultra-high field neuroimaging with the isometric handgrip paradigm, we demonstrate that arousal-driven LC activity dynamically shapes attention allocation and recognition memory for emotionally salient information. Using a novel high-resolution perfusion imaging protocol, we demonstrate that our physiological arousal manipulation via isometric handgrip selectively increases metabolic activity in the LC. Our study provides mechanistic insight into how physiological arousal shapes salience and attention network dynamics, with important implications for alterations of these mechanisms in affective disorders, aging, and neurodegenerative diseases.

## Author Contributions

AJK, IP, MJD, HILJ, and MM designed the experiment. AJK created the task code and experiment. HILJ provided the 7T MT scan parameters. IP created the task scan sequences. CZ and DJJW created the ASL scans. AJK and IP collected the task data. CZ and FG collected the ASL data. CZ and FG did the ASL scan reconstruction and ASL data curation. AJK conducted data analyses. AJK wrote the initial manuscript draft. All authors revised the manuscript.

## Funding Statement

This work is supported by the National Institutes of Health (T32AG000037 to AJK; R01AG062559, R01AG068062, R01AG082006 to HILJ; R01EB032169, R01NS134712 to DJJW), the Alzheimer’s Association (AARG-22-920434 to HILJ), and the BrightFocus Foundation (ADR-A2024006F to MJD).

## Declarations of Competing Interests

none

## Notes

### Competing Interest Statement

The authors have declared no competing interest.

### Summary of Updates

Reorganized sections so methods comes earlier.

